# Genetic analysis identifies the *Ostrea stentina/aupouria/equestris* oyster species complex in Hawai‘i, and resolves its lineage as the western Pacific *O. equestris*

**DOI:** 10.1101/2020.03.22.002444

**Authors:** Jolene T. Sutton, Jared Nishimoto, Jeremy Schrader, Keinan Agonias, Nicole Antonio, Brandi Bautista, Riley Cabarloc, Maata Fakasieiki, Noreen Aura Mae Gonong, Torey Ramangmou, Lavin Uehara, Jade Wong, Daniel Wilkie, David Littrell, Marni Rem-McGeachy, Rhiannon Chandler-‘Īao, Maria Haws

**Affiliations:** Department of Biology, University of Hawai‘i at Hilo, 200 W. Kāwili St. Hilo, Hawai‘i 96720; Tropical Conservation Biology and Environmental Science (TCBES) program, University of Hawai‘i at Hilo, 200 W. Kāwili St. Hilo, Hawai‘i 96720; Pacific Aquaculture & Coastal Resources Center (PACRC), University of Hawai‘i at Hilo, 1079 Kalanianaole Avenue, Hilo, Hawai‘i 96720; Waiwai Ola Waterkeepers Hawaiian Islands

**Author notes:** Contributed equally, and listed alphabetically by last name (Keinan Agonias, Nicole Antonio, Brandi Bautista, Riley Cabarloc, Maata Fakasieiki, Noreen Aura Mae Gonong, Torey Ramangmou, Lavin Uehara, Jade Wong). Corresponding author Corresponding Author: Jolene Sutton, 200 W. Kāwili St., Hilo, HI, 96720, U.S.A.

## Abstract

**Background:** Extensive phenotypic plasticity in oysters makes them difficult to identify based on morphology alone, but their identities can be resolved by applying genetic and genomic technologies. In this study, we collected unknown oyster specimens from Hawaiian waters for genetic identification.

**Methods:** We sequenced two partial gene fragments, mitochondrial 16S ribosomal RNA (*16S*) and cytochrome c oxidase subunit I (*COI*), in 48 samples: 27 unidentified oyster specimens collected from two locations on O‘ahu, 13 known specimens from a hatchery in Hilo, Hawai‘i Island, and 8 known specimens from Hilo Bay, Hawai‘i Island.

**Results:** Molecular data identified approximately 85% of unknown samples as belonging to the *Ostrea stentina/aupouria/equestris* species complex, a globally distributed group with a history of uncertain and controversial taxonomic status. The remaining unknown samples were the native *Dendostrea sandvichensis* (G. B. Sowerby II, 1871), and nonnative *Crassostrea gigas* (Thunberg, 1793), the latter of which is a commercial species that was introduced to Hawai‘I from multiple sources during the 20th century. Phylogenetic analysis placed Hawai‘i *Ostrea* alongside samples from China, Japan, and New Zealand, grouping them within the recently classified western Pacific *O. equestris*. Until now, four extant species of true oyster have been documented in Hawai‘i. This study expands the known range of *O. equestris* by providing the first verification of its occurrence in Hawai‘i.

## Introduction

Cryptic morphology within and among oyster species causes taxonomic confusion and may complicate aquaculture and management efforts, however, oyster identities and evolutionary histories are increasingly being resolved using genetic and genomic technologies (*e.g.* Guo et al. 2018; Hamaguchi et al. 2017; Li et al. 2017a; Li et al. 2017b). In Hawai‘i, resurgence in traditional fishpond aquaculture and an associated interest in farming both native and non-native oysters has spurred recent interests to identify unknown species. Native and non-native oyster cultures have also been applied to water quality improvement in polluted areas. These commercial and environmental efforts are complicated both by the wide phenotypic variation among oysters and by the large number of bivalve species that are present.

Hawai’i has over 1000 species of mollusk, of which approximately 160 are bivalves (Kay 1979; Moretzsohn & Kay 1995). These include at least four native true oyster species (Family Ostreidae (Moretzsohn & Kay 1995): *Dendostrea sandvichensis*, *Parahyotissa numisma* (Lamarck, 1819), *Lima lima* (Linnaeus, 1758), and *Neopycnodonte cochlear* (Poli, 1795). A number of cupped oysters are also suspected to be native (*e.g. Pinctada margaritifera* (Linnaeus, 1758)). Additional species have been introduced for commercial purposes (*e.g. C. virginica* (Gmelin, 1791), and *C. gigas)* or through possible accidental introductions (*e.g. Saccostrea sp.*). It is likely that multiple undocumented oysters exist which represent both native and introduced species. For example, at least three additional species of *Ostrea* were recorded as early as 1912 (Pilsbry 1917) which were not clearly assigned to species (Coles et al. 1999; Coles et al. 1997). During more recent surveys on O‘ahu between 2016 to 2018, we observed further specimens that did not appear to correspond to the taxonomic and shell morphology traits of species known to inhabit Hawai’i.

At present, Hawaii’s aquaculture industry is largely focused on introduced, globally commercial species for food consumption. However, the industry may benefit by increasing utilization of native species. For example, surveys show that chefs are willing to pay an additional $5.25 per dozen for oysters that are grown locally, suggesting that labeling locally-grown oysters may be a valuable marketing strategy (Chen et al. 2017b) even for introduced species. This work suggests that there may also be a market for labeling native oyster species. With the exception of *D. sandvichensis*, the commercial potential of Hawaii’s native true oysters is somewhat limited due to their small size, although *D. sandvichensis* may reach sizes of over five to six centimeters in length, similar to that of Kumamoto oysters (*C. sikamea*). If currently undocumented species can be identified and observed already growing well in Hawaiian waters, such species may present additional marketing opportunities.

Initially both aquaculture efforts and water quality efforts used non-native *C. gigas* or *C. virginica*, as these are readily available from hatcheries and the culture methods are well known (e.g. Graham et al. 2020; Grizzle et al. 2017). However, permitting issues can complicate use of introduced species, making native species an attractive option, especially as the latter may be better adapted to the wide variety of environments found in Hawai’i and/or have less invasion risk compared to non-native species. Culture of *D. sandvichensis* first took place in He‘eia fishpond (O‘ahu) and Hale O Lono fishpond (Hawai’i Island) in 2011-2013 (Thompson & Butler 2019). In terms of water quality applications, successful trials with suspension-feeding bivalves have been conducted in Hilo Bay, Kāne‘ohe Bay, Sand Island, Pearl Harbor, and at two marinas at the mouth of the Ala Wai Canal, O‘ahu. These efforts are the result of a partnership with the Waiwai Ola Waterkeepers, the University of Hawai’i at Hilo Pacific Aquaculture & Coastal Resources Center (PACRC), the U.S. Navy, U.S. Marine Corps, and the Polynesian Voyaging Society to begin utilizing native bivalves for water quality mitigation and eventual restoration of the species (Thompson & Butler 2019). Given the need for current information about the presence and distributions of native and nonnative oyster species in order to support a growing aquaculture industry in Hawai’i and for informing regulatory decisions, we used DNA barcoding (Hebert et al. 2003) to identify samples that did not correspond to the traits associated with previously documented oyster species.

## Materials & Methods

### Sample collection

We sampled a total of 27 unidentified oysters collected from two locations on the Island of O’ahu (Kāne’ohe Bay, n=3; Pearl Harbor, n=13; mixed samples sourced from Kāne’ohe Bay and Pearl Harbor, n = 11; Table 1; Figure 1). We also sampled 9 *D. sandvichensis* and 4 *C. gigas* from a hatchery in Hilo on the Island of Hawai’i, and 8 *D. sandvichensis* from Hilo Bay. We excised adductor and gill tissues from each sample and either processed them immediately or preserved them in 70% ethanol until DNA extraction. The specimens from Kāne’ohe Bay were found the He’eia fishpond, and were typically found in the upper intertidal zone attached to hard surfaces such as rocks or artificial substrates. The specimens of *D. sandvichensis* were either found in the lower intertidal zone or subtidally. *Crassostrea virginica* was observed to occur either in the same zone as the unidentified specimens, or higher in the upper intertidal zone. The distribution of *C. gigas* in Kāne’ohe Bay was variable and extended into the lower intertidal area. The unidentified specimens from Pearl Harbor were collected from the West Loch. *Crassostrea gigas* was found less frequently in Pearl Harbor compared to Kāne’ohe Bay, and was observed in the same zone as *C. virginica*.

**Table 1:**
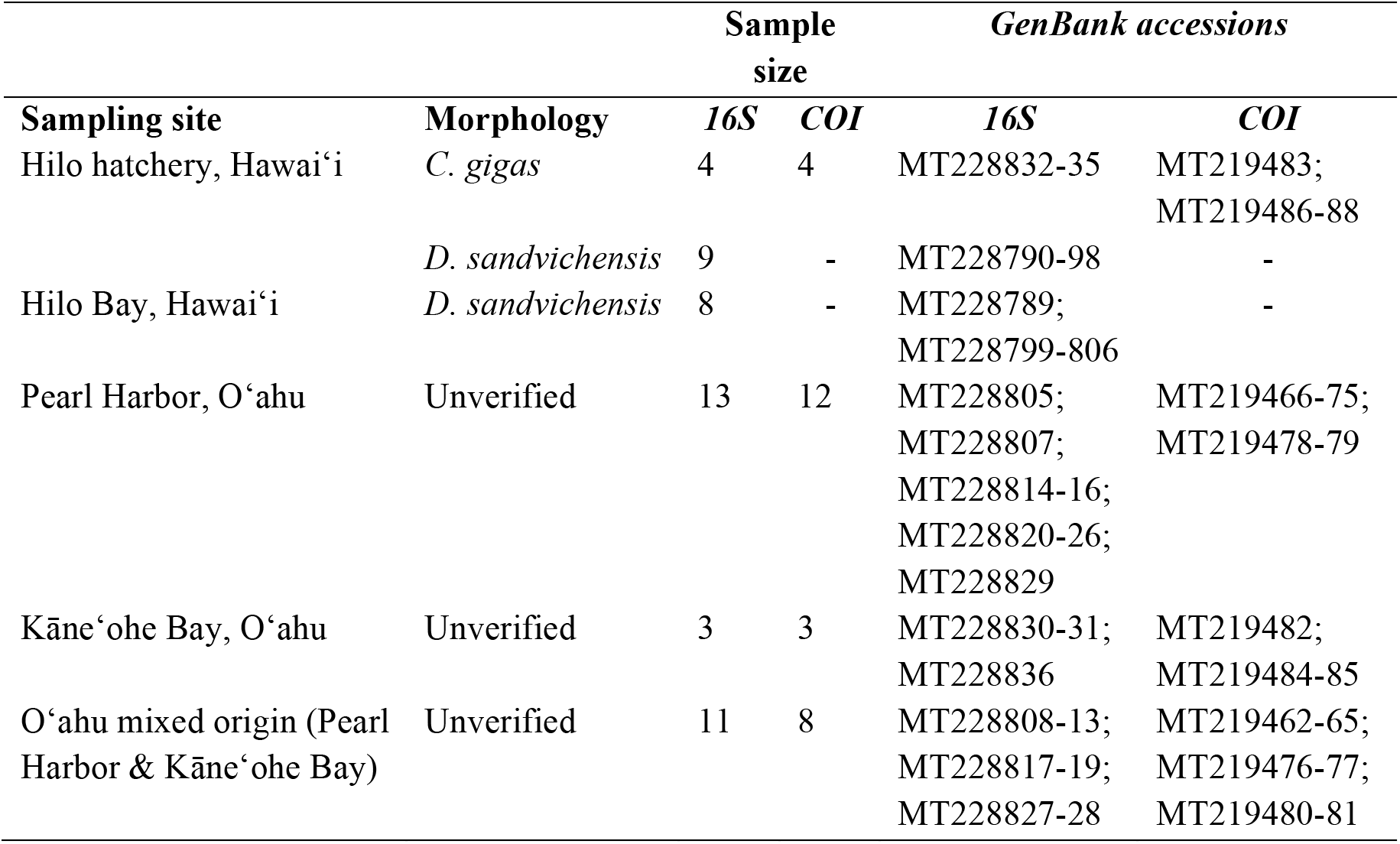
Locations, sample sizes, identities at time of collection, and GenBank sequence accession numbers for samples collected in this study.

**Figure 1:**
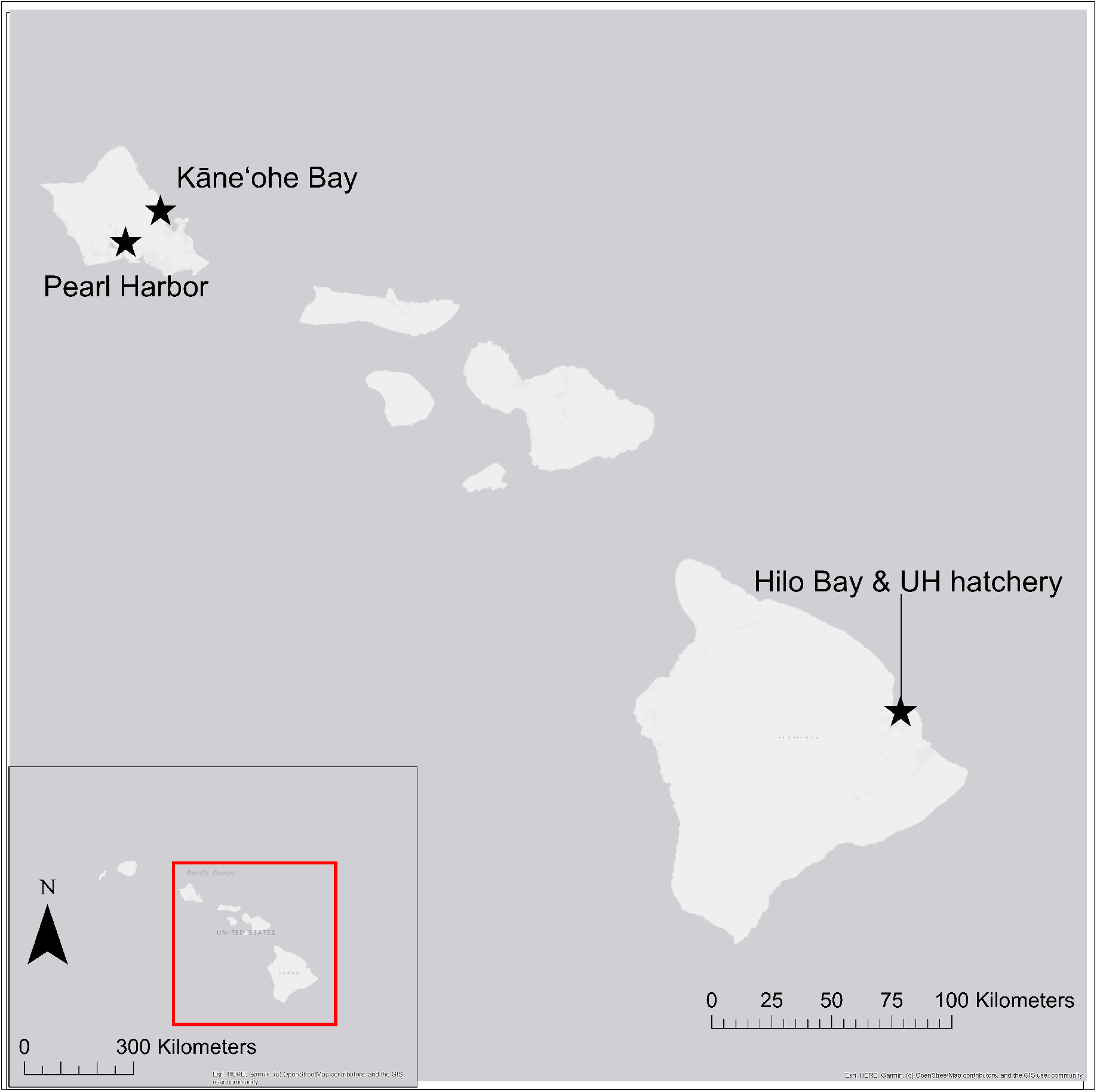
Map of the State of Hawai’i (inset) highlighting the sampling locations on O’ahu (Kāne’ohe Bay and Pearl Harbor) and Hawai’i Islands (Hilo Bay and the University of Hawaii at Hilo hatchery.

### Molecular methods and BLASTn searches

We used a DNeasy Blood and Tissue Kit (Qiagen) to extract gDNA from each tissue sample, and we assessed DNA quality and quantity via gel electrophoresis (1.5% agarose in TAE stained with GelRed® [Biotium]) alongside a 10 kb ladder (Thermo Fisher Scientific GeneRuler™ DNA Ladder Mix #SM0334). We standardized DNA concentrations of high molecular weight samples to ~20ng/*μ*l prior to PCR. If DNA appeared sheared, we did not adjust concentrations. To amplify partial gene regions of *COI* and *16S*, we used standard barcoding primers LCO1490 and HC02198 (*COI*; Folmer et al. 1994), and 16Sar-L and 16Sbr-H (*16s*; Palumbi et al. 1991). For each gene, we prepared PCRs in final 25 *μ*l volumes consisting of 1× buffer (NEB Hot Start Taq® 2X Master Mix, catalogue #M0496; final concentrations: 25 units/ml *Taq* DNA polymerase, 1.5 mM MgCl_2_ and 0.2 mM each dNTP), 0.2 *μ*M of each *COI* primer or 0.4 *μ*M of each *16S* primer, and ~20 ng of DNA. Our thermal cycling conditions for *COI* consisted of initial denaturation at 95 °C for 45 s followed by 10 touchdown cycles of 95 °C for 20 s, Ta for 40 s, 68 °C for 60 s. This was followed by 30 cycles of 95 °C for 20 s, 40 °C for 40 s, 68 °C for 60 s and a final extension of 68 °C for 5 minutes. During the touchdown cycles, the starting annealing temperature was 50 °C, and each cycle reduced the annealing temperature by 1 °C. For *16S*, we relied on initial denaturation at 95 °C for 30 s followed by 12 touchdown cycles of 95 °C for 30 s, Ta for 60 s, 68 °C for 60 s. This was followed by 25 cycles of 95 °C for 20 s, 40 °C for 40 s, 68 °C for 60 s and a final extension of 68 °C for 5 minutes. During the touchdown cycles, the starting annealing temperature was 55 °C, and each cycle reduced the annealing temperature by 1 °C. We assessed amplification success and checked negative controls via gel electrophoresis as previously described, and we used ExoSAP-IT™ (Thermo Fisher Scientific) to purify the PCR products. We Sanger sequenced each sample in forward and reverse directions at the University of Hawai’i at Mānoa Advanced Studies in Genomics, Proteomics and Bioinformatics (ASGPB) facility, and we used Geneious (Biomatters 2019) to view and edit sequences and for subsequent phylogenetic analysis.

We used NCBI’s BLASTn search to compare our sequences against known references on GenBank (Sayers et al. 2018). Our BLASTn searches identified the known *D. sandvichensis* samples as *D*. *crenulifera* (G. B. Sowerby II, 1871), which is a synonym for *sandvichensis* (WoRMS 2020). Similarly, our known *C. gigas* were verified as *C. gigas*. Of our 27 unknown samples, we generated *16S* sequences for all 27, and we generated *COI* sequences for 23 samples. Using *16S*, 23/27 unverified samples were identified as belonging to the *O. stentina/aupouria/equestris* species complex, 3/27 were identified as *C. gigas*, and 1/27 was identified as *D. sandvichensis*. Using *COI*, 20/23 unverified samples were identified as belonging to the *O. stentina/aupouria/equestris* species complex, and 3/23 were identified as *C. gigas* (Table 2). GenBank accessions for sequences generated in this study are MT228789-MT228836 (*16S*) and MT219462-MT219488 (*COI*).

**Table 2:** Sources and genetic identities of samples that were unverified from morphology at the time of collection. The *COI* locus only includes data for *Ostrea* and *Crassostrea*, due to inconsistent amplification of *Dendostrea*.

| Genetic identification | Pearl Harbor | Kāne‘ohe Bay | O‘ahu mixed |
| --- | --- | --- | --- |
| **16S (n = 27)** |  |  |  |
| *O. stentina* complex | 12 | 0 | 11 |
| *C. gigas* | 0 | 3 | 0 |
| *D. sandvichensis* | 1 | 0 | 0 |
| **COI (n=23)** |  |  |  |
| *O. stentina* complex | 12 | 0 | 8 |
| *C. gigas* | 0 | 3 | 0 |

### Phylogenetic analysis

After trimming sequences to including only the inter-primer region, we used ClustalW (Thompson et al. 2003; Thompson et al. 1994) implemented in Geneious to align our sequences with references from GenBank. We used a Bayesian inference (BI) approach for separate phylogenetic analyses of *16S* and *COI* sequences to examine taxonomic separation across species (*Ostrea*, *C. gigas*, and *D. sandvichensis*). To select the best fitting models for BI analysis, we used jModelTest v2.1.10 (Darriba et al. 2012; using default parameters and the three substitution scheme for analyzing models that could be implemented in MrBayes (Huelsenbeck & Ronquist 2001)) and the Bayesian information criterion (BIC). The selected models were HKY+I+G and HKY+G respectively for *16S* and *COI*. We constructed Bayesian trees via MrBayes implemented in Geneious, using the following species for comparison: *C. gigas* (DQ839414, FJ743509, AY632550, JF808180, JF700177 (Pie et al. 2006; Sayers et al. 2018; Wang et al. 2004; Zhang et al. 2013)), *C. gigas angulata* (KC170323, KC170322 (Peng-yun 2013; Sayers et al. 2018)), *D. sandvichensis* (KC847121, EU815985 (Sayers et al. 2018)), *O. angasi* (AF052063 (Jozefowicz & Foighil 1998; Sayers et al. 2018)), *O. angelica* (KT317127, KT317140, KT317449 (Raith et al. 2015; Sayers et al. 2018))*, O. chilensis* (AF052065 (Jozefowicz & Foighil 1998; Sayers et al. 2018)), *O. circumpicta* (MG560202 (Sayers et al. 2018), *O. conchaphila* (KT317173, FJ768528, KT317494 (Polson et al. 2009; Raith et al. 2015; Sayers et al. 2018)), *O. denselamellosa* (FJ743511, HQ660995, KP067907 (Kim et al. 2015; Liu et al. 2011; Sayers et al. 2018)), *O. edulis* (JQ611449, AF540595, KJ818235 (Malkowsky & Klussmann-Kolb 2012; Morton et al. 2003; Pejovic et al. 2016; Sayers et al. 2018)), *O. futamiensis* (LC051603 (Hamaguchi et al. 2017; Sayers et al. 2018)), *O. lurida* (FJ768559, FJ768554, KT317504 (Polson et al. 2009; Raith et al. 2015; Sayers et al. 2018)), *O. permollis* (AY376605, AY376606, DQ226526 (Kirkendale et al. 2004; Sayers et al. 2018))*, O. puelchana* (AF052073, DQ226521 (Jozefowicz & Foighil 1998; Sayers et al. 2018)), and *O. stentina/aupouria/equestris* species complex (Table 3). For outgroups, we used sequences from *Hyotissa imbricata* (KC847136 and AB076917 (Matsumoto 2003; Sayers et al. 2018)). We set our MrBayes parameters were as follows: total chain length = 1,100,000; burnin = 100,000; sub-sampling frequency = every 200 iterations. We ran four heated chains per analysis. To confirm model convergence, we examined the posterior outputs of each statistic for the minimum effective sample size (minESS; convergence based on minESS ≥ 200), the posterior scale reduction factor (PSRF; convergence based on PSRP = 1), and made visual checks of the trace and density graphs.

**Table 3:**
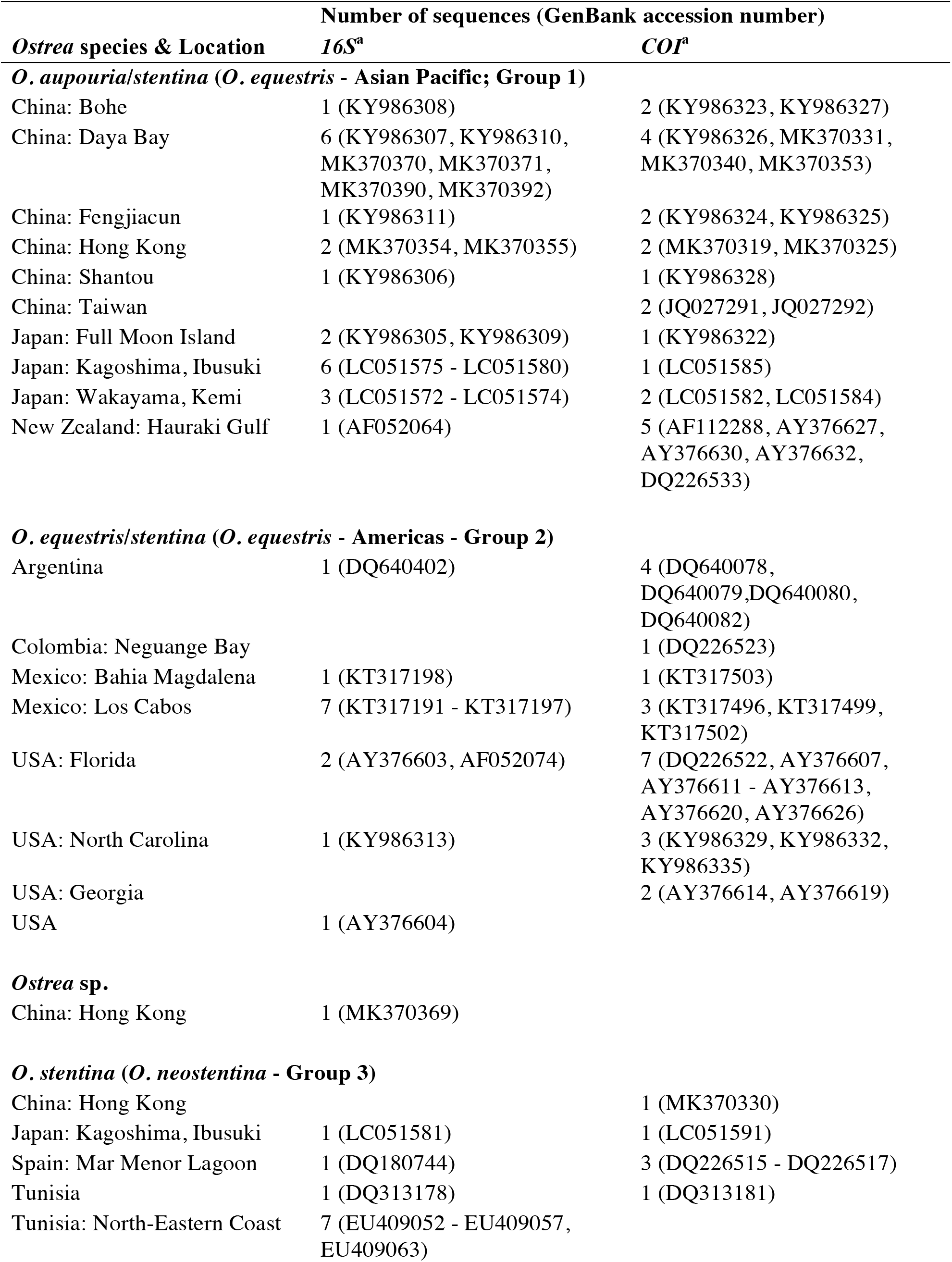
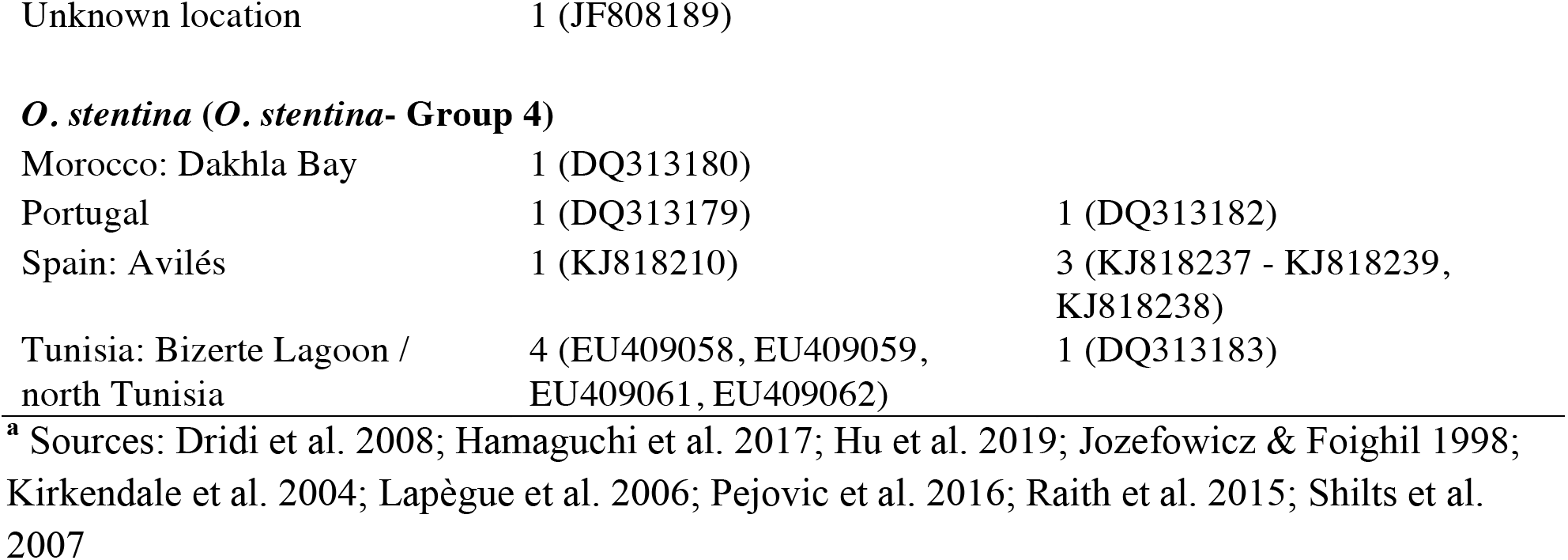
Locations and sequence accession numbers of *Ostrea stentina* species complex samples from GenBank. Parentheses indicate the species classification of Hu *et al.* (2019).

## Results

### Analysis of 16S

From 48 oyster specimens collected and identified in this study, we obtained 48 *16S* and 27 *COI* sequences. The *16S* analysis comparing our sequences with representative *C. gigas, D. sandvichensis*, and *Ostrea* sequences from GenBank confirmed the identities of our known samples, and resolved the identities and relationships of our unknown samples (Figure 2). All of our *C. gigas* and *D. sandvichensis* samples clustered with strong support (PP=1 and PP=0.998) alongside representative sequences from GenBank. Similarly, our *Ostrea* samples fell into a well supported clade (PP = 1) alongside specimens from the *O. stentina/aupouria/equestris* species complex, which were recently recommended as a separate species, *O. equestris* (Hu *et al.* 2019). Two strongly supported clades (PP = 0.998 and 1) that excluded Hawai’i *Ostrea* distinguished groups 3, and 4 from Hu *et al.* (2019), representing *O. neostentina*, and *O. stentina*, respectively. Of 13 *O. equestris* (Americas) sequences, 10 formed a moderately supported clade (PP = 0.868). There was no power to differentiate *C. gigas* samples by origin (Figure 2). There was evidence to suggest some differentiation among *D. sandvichensis*, with a well supported clade (PP = 0.960) comprised of nine hatchery samples and one Hilo Bay sample. Excluding the outgroup, the overall average evolutionary distance (measured as the sum of branch lengths) was 0.074 and ranged from 0.002 to 0.255. The greatest distance was between *O. angasi* (AF052063) and *C. gigas angulata* (KC170322). The smallest difference between Hawai’i *Ostrea* and GenBank samples (0.002) occurred with two samples from China (LC051572 and LC051574).

**Figure 2:**
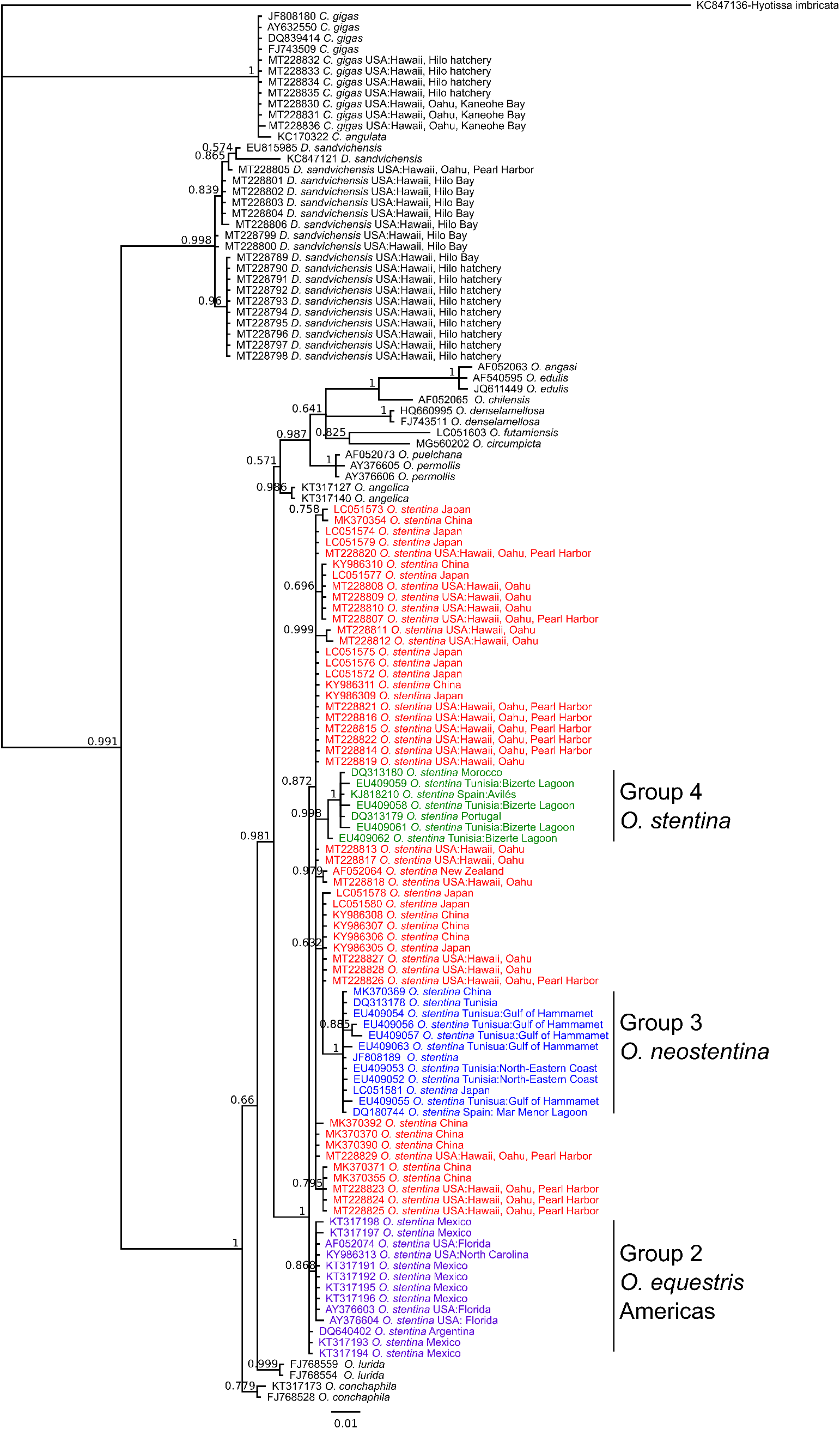
Bayesian inference tree of oyster species based on 341-488 bp of *16S*. Bayes posterior probabilities (PP) greater than 0.5 are given for each node. Accession numbers are used for samples obtained from GenBank. Group designations match those of Hu *et al.* (2019): Group 1 (red) = *O. equestris* (western Pacific); Group 2 (purple) = *O. equestris* (Americas); Group 3 (blue) = *O. neostentina*; and Group 4 (green) = *O. stentina*.

### Analysis of COI

Samples identified as *Dendostrea* from our *16S* analysis inconsistently amplified from the standard *COI* primers, and were therefore removed from our analysis of this locus. Our remaining known samples were confirmed as *C. gigas*, and our unknown samples were resolved as either *C. gigas* or *O. stentina* in accordance with their identities based on *16S* (Figure 3). There was no power to differentiate samples by origin for *C. gigas* (Figure 3). All of our *Ostrea* samples clustered with representative samples of the *O. stentina/aupouria/equestris* complex. Moderately to strongly supported clades distinguished groups 2, 3, and 4 from Hu *et al.* (2019) and excluded Hawai’i specimens (PP = 0.796, 0.988, and 1 respectively). All Hawai’i samples grouped in a strongly supported clade (PP=0.941) together with group 1, *O. equestris* (western Pacific), from Hu *et al.* (2019). Excluding the outgroup, the overall average evolutionary distance (measured as the sum of branch lengths) was 0.097 and ranged from 0.002 to 0.399. The greatest distance was between a *C. gigas* from Hawai’i (MT228832) and *O edulis* (KJ818235). The smallest difference between Hawai’i *Ostrea* and GenBank samples (0.002) occurred with samples from China (KY986323, MK370325, JQ027291), Japan (KY986322, LC051582), and New Zealand (AY376627).

**Figure 3:**
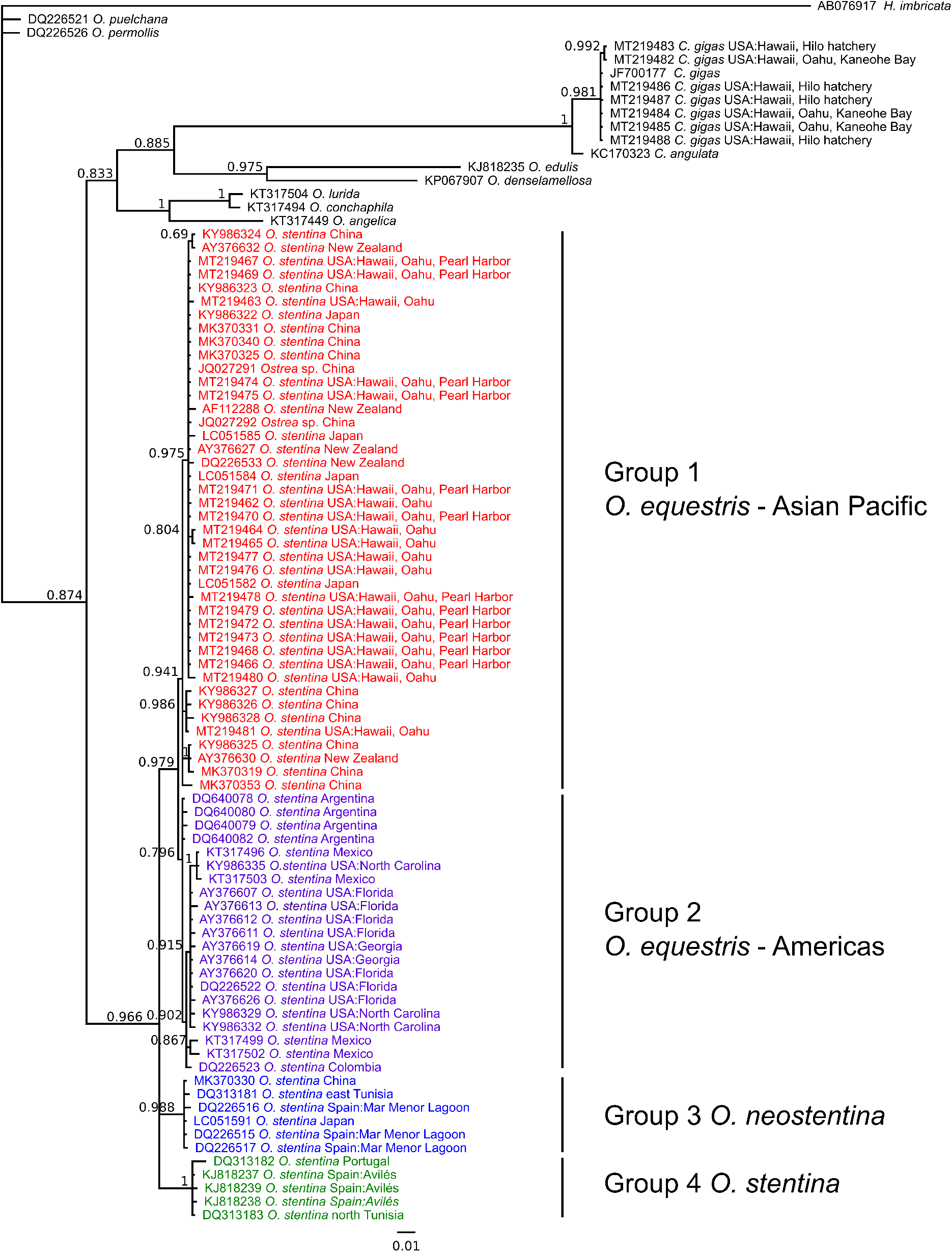
Bayesian inference tree of oyster species based on 453-651 bp of *COI*. Bayes posterior probabilities (PP) greater than 0.5 are given for each node. Accession numbers are used for samples obtained from GenBank. Group designations match those of Hu *et al.* (2019): Group 1 (red) = *O. equestris* (western Pacific); Group 2 (purple) = *O. equestris* (Americas); Group 3 (blue) = *O. neostentina*; and Group 4 (green) = *O. stentina*.

### Morphology of Hawai’i Ostrea

The shell shape for samples genetically identified as *Ostrea* were typical of the genus, being generally oval, although some specimens were elongated or nearly triagonal. In the latter cases, an inwardly curving indentation along the margin sometimes occurred, giving the shells a somewhat heart-shaped appearance. The left valve was sometimes slightly larger and more cupped than the right. The exterior shell color was whitish gray, with a greenish tinge in most specimens. The shell was not rayed and was often worn, revealing slightly greenish shell layers. The interior shell color was usually white or white tinged with green. The nacreous layer had more orient than in the other species of *Ostrea* or *Crassostrea* found in Hawai’i, giving it a luminescent quality. The adductor muscle scar ranged from inconspicuous to a light brown color. Minor chromata were found on the inside of the shell on the margin in some specimens. More conspicuous chromata were usually confined to areas near the hinge, although a few specimens had barely noticeable chromata around the entire shell margin. The hinge ligament was external and alivincular, and not pronounced. No teeth were present. The largest dorsoventral measurement (DVM) was 60.3 mm.

## Discussion

This study expanded the known distribution of *O. equestris* by confirming its presence in Hawai’i. It also provided evidence to suggest there may be genetic structuring in the native *D. sandvichensis*, based on differences between samples from the University of Hawai’i at Hilo hatchery and from the wild. This is the first documented record of *O. equestris* in Hawaiian waters. The *O. stentina/aupouria/equestris* species complex is globally distributed, found along Atlantic coasts, the Mediterranean, North Africa, New Zealand, Japan, China, and South America (Crocetta et al. 2013; Hamaguchi et al. 2017; Hu et al. 2019; Lapègue et al. 2006; Pejovic et al. 2016). At this time, it is uncertain whether *O. equestris* in Hawai’i is native or introduced, though it seems likely to be native given the wide distribution of *O. equestris* in both the Atlantic and Pacific oceans. Further, *Ostrea* traits such as hermaphroditism and small size have been suggested to be advantageous for range expansion on equatorial currents of the Pacific (Hu et al. 2019).

A recent classification of the *Ostrea* genus identified four closely related groups in the *O. stentina/aupouria/equestris* species complex, and recommended they be designated as separate species (Hu et al. 2019). These were group 1: *O. aupouria/stentina* from China, Japan, and New Zealand, which was recommended as *O. equestris* (western Pacific); group 2: *O. equestris* from Gulf of California, Argentina, Florida, and North Carolina, which was recommended as *O. equestris* (Americas); group 3: *O. stentina* from southeastern Spain and eastern Tunisia, which Hu *et al.* (2019) described as a new species, *O. neostentina*; and group 4: *O. stentina* from northern Spain, Portugal, Morocco, and northern Tunisia. Guo *et al.* (2018) also suggested that the *O. stentina/aupouria/equestris* complex be distinguished as three separate species, one of which was *O. equestris*. Our phylogenetic analyses placed Hawai’i specimens with the western Pacific *O. equestris*. As with previous studies, our results support recent and ongoing speciation.

### Economic value & environmental considerations

*Ostrea equestris* is generally considered to be a small, economically unimportant species that inhabits subtidal and intertidal waters with high salinities (*e.g.* > 20%; reviewed in Galtsoff & Merrill 1962; Dinamani & Beu 1981; Markwith 2010). Its first description in New Zealand, as *O. aupouria*, measured it at 35-45mm in height (Dinamani & Beu 1981). The principal commercial *Ostrea* species (Arakawa 1990) are *O. edulis* (European flat oyster; Linnaeus, 1758), *O. lurida* (Olympia oyster, Carpenter, 1964), *O. chilensis* (Chilean oyster; Küster, 1844), and *O. angasi* (Australian flat oyster; Sowerby 1871). *Ostrea edulis* has been harvested for food since the middle part of the Stone Age, and cultivated since during the Roman Empire (Helmer et al. 2019). *Ostrea lurida*, native to the west coast of temperate western North America (Raith et al. 2015), was harvested by Native Americans for at least 4000 years, and was commercially exploited from the 1850s until 1915 (White et al. 2009). *Ostrea chilensis* is native to Chile and New Zealand, and began to be dredged in the Foveaux Strait, New Zealand, since c.1867 (Cranfield et al. 1999). *Ostrea angasi* is endemic to southern Australia, where it was originally consumed by Aboriginals and made up a commercial fishery in the 19th and 20th centuries (Alleway & Connell 2015). *Ostrea* species may be noted and prized for their unique flavor, which is often described as metallic (*e.g.* White et al. 2009).

Oyster stocks worldwide have been depleted through a combination of overexploitation, disease, pollution and invasive species (Alleway & Connell 2015; Helmer et al. 2019; Lotze et al. 2006). Natural beds of *O. edulis*, for example, were intensely harvested from the 18th century until the middle of the 19th century when overexploitation led to reduced stocks and fishery restrictions (Buestel et al. 2009; da Silva et al. 2005; Helmer et al. 2019). Several *Ostrea* species are susceptible to the parasites *Bonamia exitiosa* and *Bonamia sp.*, which have also impacted food fisheries of *O. edulis* (Helmer et al. 2019) and *O. chilensis* (Berthe & Hine 2003; Cranfield et al. 2005). *Bonamia* infections have also been found in *O. stentina* in Tunisia (Hill et al. 2010) and the U.S.A (Hill et al. 2014; note this is likely to be *O. equestris* based on the classification of Hu *et al.* [2019]), *O. angasi* (Corbeil et al. 2006), and *O. lurida* (Engelsma et al. 2014), though not all species appear susceptible (Engelsma et al. 2014). *Bonamia* have also been identified in non-commercial species such as *O. equestris* (Engelsma et al. 2014), and in *D. sandvichensis* from Hawai’i (Hill et al. 2014), among others. In addition to providing a food resource, oysters are valued for ecosystem services, and current restoration and cultivation efforts aim to replenish depleted food fisheries as well as improve damaged marine environments. Services provided by oysters notably include water filtration which reduces eutrophication (Alleway & Connell 2015; Zu Ermgassen et al. 2013), and habitat engineering (Beck et al. 2011) which increases habitat complexity along with diversity of fish and other species (Alleway & Connell 2015; Helmer et al. 2019). Degradation of oyster reefs has negatively impacted ecosystem functions worldwide (Alleway & Connell 2015; Beck et al. 2011; Helmer et al. 2019).

### Hawai’i aquaculture

Native Hawaiians practiced aquaculture for centuries prior to the arrival of Europeans, however, the local fishpond productivity was reduced 100-fold by the mid 1970s (Chen et al. 2017a). In recent years, the industry has begun to expand once more, and to diversify by producing oysters for environmental purposes as well as for food. The documentation of *O. equestris* in Hawai’i offers a potential opportunity to contribute to these efforts, especially if the species is native. For example, one area into which the industry has expanded is using suspension-feeding bivalves to improve water quality. The use of native species is an attractive potential option for these purposes. Along with *D. sandvichensis*, locally produced *O. equestris* holds potential to differentiate Hawaii’s shellfish farms. The observation of this species growing well in Hawaiian fishponds may also offer an option for production in those systems. Use of native species could synergize with commercial efforts while helping to restore native bivalve stocks.

## Conclusions

By documenting *O. equestris* in Pearl Harbor, we expanded the known distribution of this species. We recommend that additional surveys be conducted to assess the distribution and habitat requirements of *O. equestris* throughout Hawai’i. Future studies could also assess disease susceptibility of this species, as well as develop protocols for its cultivation. Genomic research could better assess its evolutionary history and origins.

## Acknowledgements

Thanks to Debbie Beirne for maintaining the Biology teaching laboratories at the University of Hawai’i at Hilo. Thank you to Erin Datlof and Hallie Whitmore for assistance in the molecular laboratory. Thank you to Matthew Knope for helpful feedback on the phylogenetic analysis. We also thank the following entities for allowing the collection of specimens and for other support: U.S. Navy team at the Pearl Harbor Joint Base Hickam, Paepae O He’eia and the Hawai’i Institute of Marine Biology.

